# Ancient environmental DNA indicates limited human impact on marine biodiversity in pre-industrial Iceland

**DOI:** 10.1101/2024.09.29.615643

**Authors:** Luke E. Holman, Emilia M. R. Arfaoui, Lene Bruhn Pedersen, Wesley R Farnsworth, Phillipa Ascough, Paul Butler, Esther R. Guðmundsdóttir, David J. Reynolds, Tamara Trofimova, Jack T. R. Wilkin, Christian Carøe, Tobias Guldberg Frøslev, Ramona Harrison, Shyam Gopalakrishnan, Mikkel Winther Pedersen, James Scourse, Kristine Bohmann

## Abstract

Human activities are affecting marine biodiversity globally by accelerating extinction rates, altering ecosystem conditions, and changing community structures. These changes can only be understood through establishing the ecosystem state prior to significant anthropogenic impact, and by disentangling the anthropogenic effect from natural climatic changes. Here, we reconstruct marine biodiversity in Iceland across three millennia (1315 BCE-1785 CE), encompassing periods of climatic fluctuation and human settlement, to explore the comparative effect of natural and anthropogenic forces on marine biodiversity. We performed 18S metabarcoding of ancient environmental DNA from two sediment cores collected from northern Icelandic shelf seas, integrating local climatic records, population estimates and zooarchaeological remains from published sources to estimate the influence of climatic and anthropogenic impacts. Against the backdrop of increasing human populations and marine exploitation, we observe no large-scale taxonomic shifts or anthropogenic biodiversity changes across the period. In contrast, we found a positive correlation between herring (*Clupea harengus*) detection rates and proxy-reconstructed sea surface temperature, suggesting a role for climate in shaping marine biodiversity. Overall, our data suggest that despite impacts on terrestrial ecosystems and the development of a substantial export fishery across the study period, Icelandic society may have had a limited effect on marine biodiversity.

## Introduction

Some of the most abrupt global marine ecosystem changes have occurred across recent decades [1,2], with marine biodiversity under threat from a suite of anthropogenic stressors [3]. Biodiversity time-series data can help to document the scale and rate of this change, which is important to improve our understanding of ecology, but is also key in marine resource management, to minimise the loss of ecosystem services [1], and to set goals for ecosystem restoration [2]. However, most datasets span only a few decades, and even the longest continuous surveys provide little information about pre-industrial marine ecosystems [4]. This is critical as there is a growing awareness of the hitherto unseen impact of prehistoric and preindustrial human societies on ecosystems [5,6]. Research from historical ecology and archaeology has revealed the size and breadth of human impact, well beyond the timescale of modern monitoring [5,6]. Data from these disciplines provides a view of marine anthropogenic impact from a human perspective, analysing data associated with human sites or records, for example fisheries catch logs or bone material from kitchen middens. However, they rarely measure biodiversity change in the ecosystem directly. In contrast, palaeoecology provides data on natural ecosystems across vast ecological (and evolutionary) time, but only a limited fraction of biodiversity is fossilised and therefore preserved for analysis [7]. Together, this means we have limited information about past marine biodiversity against which to compare modern biodiversity change. This is further complicated because ecosystem conditions within living memory are assumed to be “normal” while longer term changes remain unrecognised; so-called shifting baseline syndrome [8]. Collectively this means we lack both a comprehensive understanding of baseline pre-human biodiversity, and also a full accounting of the effect of humans on marine ecosystems.

Throughout their lifetime organisms shed DNA into the environment; this genetic material can be deposited and incorporated into marine sediments, which accumulated over time preserves an archive of past biodiversity. Analysis of this ancient environmental DNA (eDNA) is an approach that is becoming increasingly common to study past marine ecosystems [9]. The analysis of marine sediment archives has enabled the reconstruction of past fish abundances [10], revealed marine ecosystem change as a result of both natural [11] and anthropogenic [12] forcings, and reconstructed previously undescribed marine ecosystems from millions of years ago [11,13]. The two most commonly used DNA analysis approaches to describe biodiversity data from marine eDNA data are metagenomics and metabarcoding. While both generate high-throughput sequencing data, metabarcoding provides data on taxonomically informative DNA markers for a particular taxonomic group, while shotgun-metagenomics provides more complex genome-wide data. Research indicates that across the last < 3-5k years, when damage is unlikely to have accumulated, both metabarcoding and metagenomics produce reliable biodiversity data from marine sediments [14]. However, in contrast to metabarcoding, metagenomics allows authentication of detections by measuring DNA-damage through time [11,13,14], but is currently less methodologically developed than metabarcoding, has poorer DNA reference database coverage and fewer established bioinformatic methods [15].

Iceland serves as an ideal case study to evaluate the effect of human impacts on marine biodiversity with ancient eDNA. Located just south of the Arctic Circle and surrounded by the inhospitable North Atlantic Ocean, Iceland represented a remote unclaimed island for early seafarers and was one of the last areas of the world to be settled [16]. The earliest permanent settlement sites in Iceland are associated with the Landnám tephra (expelled volcanic material) sequence dated to 877 CE, with settlers of Norse and Gaelic genetic ancestry [16,17]. The Icelandic legislative assembly (Alþingi) was the dominant power in the commonwealth period from 930-1262 CE, this was followed by rule under Norwegian and Danish kings from 1262 to 1944 CE, before the founding of the Republic of Iceland in 1944 CE [18]. Evidence from lake core sedimentary ancient DNA indicates that terrestrial plant species composition remained relatively stable from 10.0 - 1.0 cal. kyr BP [19], suggesting the Icelandic settlers would have found a pristine sub-arctic terrestrial environment following biotic recolonisation after Icelandic deglaciation during the late Pleistocene (15.4 - 14.6 cal. kyr BP) [20]. Marine palaeoecological data from northern Iceland reveals ecological responses to shifts in Holocene climate [21,22], indicating post-glacial establishment of shelf sea ecosystems, which would also likely have been pristine at the advent of settlement.

The impact of early Icelanders on the environment was largely determined by the subsistence culture from which they originated. Like the Norse and Gaelic cultures during the settlement period they raised livestock and grew barley, but the northerly climate reduced yields and increased their reliance on marine natural resources, for example birds, marine mammals and fish [18,23–25]. Deforestation and grazing led to large-scale soil erosion and turnover of vegetation communities [19], resulting in a complete shift in terrestrial biodiversity shortly after settlement in some areas [but see 18]. This intensive terrestrial degradation is reflected in evidence from inland and coastal archaeological sites, which reveal a substantial majority of faunal remains from terrestrial organisms from 9-11th century followed by a shift towards predominantly marine finds from sites dating from the 11th century onwards [26]. Though only representing data from one region, stable isotope analyses of Icelandic human bone remains seem to confirm a terrestrial diet in a site (Skeljastaðir) dated to 1000–1104 CE and increased marine consumption at another site (Skriðuklaustur) dated 1493–1554 CE [27]. The bulk of marine archeo-faunal remains from Icelandic sites referred to here are from gadid species (family Gadidae), with Atlantic cod (*Gadus morhua*), hereafter cod, being dominant (>80% in the majority of assemblages in [26]). Besides being a focus for subsistence, cod was also the basis of an extensive and well developed fishery supplying the growing urban centres across Europe from the 13-17th centuries [26]. The Icelandic cod fishery experienced declines and failures between the 17 and 19th centuries with climate and socio-economic factors both suggested as possible explanations [26,28]. In contrast to European fisheries across the Viking and Medieval periods, Atlantic Herring (*Clupea harengus*) does not appear to have been a focus of either subsistence or export fisheries in Iceland [26,29], with substantial fisheries, and subsequent collapse, only documented in the 20th century [30,31].

Despite evidence for widespread Icelandic reliance on marine resources and large-scale export fisheries, we have only limited knowledge on the impact of the people of pre-industrial Iceland on marine environments. The only documented marine species that has been driven to extinction on Iceland during pre-industrial times is the walrus [32]. Ancient DNA analyses reveal a unique Icelandic genetic lineage of walruses [32], strongly suggesting complete human-driven walrus extirpation in the 11-12th centuries. In contrast, data from ancient cod DNA does not provide clear evidence for human fishing impacting on cod population sizes relative to larger climate-driven changes [33,34]. Moreover, stable-isotope [35] and otolith growth rate analyses [36] suggest stability of cod trophic position and growth rates over the last millenia, indicative of negligible detectable effects of humans on the most frequently fished species.

Here, we reconstruct marine biodiversity changes over three millennia from two sediment records sampled from the north Icelandic shelf. Through examining biodiversity derived from metabarcoding of ancient eDNA, in combination with previously published human population and climate data, we explore the comparative influence of Icelandic human occupation and natural climatic changes on marine biodiversity.

## Methods

### a) Sediment records

Two sediment core records from the north Icelandic shelf were sampled for ancient environmental DNA analysis: a piston core collected in May 2022 during the DY150 cruise [37] on the RRS *Discovery* (DY150-NIS-A-PC019, location: 66.551721 -17.700281), and a gravity core collected in June 2006 using a gravity corer during the Millenium B05-2006 cruise on the RV *Bjarni Sæmundsson* (B05-2006-GC01, location:, 66.5015 -19.50567). The two cores are hereafter referred to as PC19 and GC01, respectively. Both cores were split and stored at 4-5°C until sampling. Subsampling of sediments from the intact split cores followed routine ancient DNA precautions to prevent contamination [38]. These included subsampling in a laboratory with no history of PCR amplification, use of single-use full-body protective equipment, thorough cleaning with 5% bleach solution followed by 70% ethanol prior to sampling, and subsampling from the oldest to youngest sampled layer. For each sampling point, 5-10 mm of exposed core material was removed from the surface with a sterile scalpel and 5-10g of material was subsampled from the freshly exposed area using a sterile disposable 5 mL syringe. Subsamples were stored at -20 °C until DNA extraction. A total of 158 samples were taken from PC19, with sampling every 2cm from 23cm - 275cm from the core top, and additional samples taken every cm, 10cm above and below two putative tephra layers. A total of 46 samples were taken from the GC01 core, with sampling every 4 cm from the uppermost layer to 180 cm from the core top.

### b) Chronology

Eleven samples were subject to radiocarbon dating, seven for PC19 and three for GC01. For each analysed horizon, samples of mixed assemblage calcareous benthic foraminifera (range: 4-12 µg) were picked from a >125 µm sediment fraction. For four out of seven PC19 samples it was necessary to combine benthic foraminifera from layers covering 2cm to have enough material to generate precise dates; all other samples were taken from a single 1 cm layer. The midpoint of the sampling depth was used in subsequent analyses. The samples were analysed at the UK National Environmental Isotope Facility Radiocarbon Laboratory, SUERC East Kilbride, University of Glasgow. Samples were pretreated by etching with a calculated volume of 0.2 M HCl to remove the outer 20% by weight of the sample surface. Following etching, the remaining sample was completely hydrolyzed to CO_2_ using an excess of 2 M HCl. The evolved CO_2_ was cryogenically purified before conversion to graphite, and subsequently measured via Accelerator Mass Spectrometry [39].

The PC19 core chronology was aided by the identification of one tephra marker layer previously described from the shelf area north of Iceland [40–42]. Two tephra samples were extracted from visible horizons identified within PC19 at 109-110 cm and 450-450.5 cm. Samples were cleaned for humic material by wet sieving bulk sediments at 63 and 125 μm. Material was then dried, mounted in epoxy, polished and carbon-coated for geochemical analysis. Tephra geochemistry was analysed using a JXA-8230 Electron Probe Micro-Analyzer (JEOL Ltd., Tokyo, Japan) at the University of Iceland. The acceleration voltage was 15 kV, the beam current 10 nA, with a beam diameter of 5-10 μm. The standards A99 (for basaltic tephra), and Lipari Obsidian (both for silicic and intermediate tephra), were measured prior to, and after, the analyses to verify consistency in analytical conditions. Data were subsequently inspected for, and cleaned of, anomalies, specifically analyses with sums < 95% and > 101%.

A Bayesian age-depth model was generated in R (v.4.2.3) using the package Bacon (v3.2.0) [43] combining the ^14^C and tephra dates. ^14^C dates were calibrated using the MARINE20 curve [44] and all dates had a regional offset (ΔR) of 200 years ± 50 (s.d.) added to compensate for the documented reservoir effect during this period [41]. The mean date from the output of the Bayesian model was used in subsequent analyses for each horizon.

### c) DNA extraction & Metabarcoding

Pre-PCR sample processing was conducted in specialised ancient DNA facilities at the University of Copenhagen following routine ancient DNA standards to prevent contamination [38]. DNA from 0.4-0.5g of subsampled sediment from each of the 158 samples from PC19 and 46 samples from GC01 were extracted using the Qiagen MagAttract PowerSoil Pro® DNA Kit following the manufacturer’s protocol (October 2022) with minor modifications as outlined in [14]. DNA was eluted in 80 μL C6 solution. A negative extraction control was added for each batch of 30 extractions.

DNA extracts were screened for PCR inhibition by spiking them into a well characterised probe-based quantitative PCR (qPCR) targeting an artificial DNA template. The qPCR consisted of 0.4 μM forward primer (CCTCCGGCCCCTGAATG), 0.4 μM reverse primer (ACCGGATGGCCAATCCAA), 0.1 μM IDT (Integrated DNA Technologies Inc, Coralville, IA, USA) doubled-quenched probe (5’ (6-FAM)CGGAACCGA(ZEN) CTACTTTGGGTGTCCGT(IBFQ) 3’), 1x Luna Universal qPCR Master Mix (New England Biolabs, Ipswitch, MA USA), 4 μL eDNA template and 2 μL artificial target sequence (CCTCCGGCCCCTGAATGCGGCTTAGTGTGTCGTAATGGGCAACTCTGCAGCGGAACCGACTACTTTGGGTGTCCGTGTTTCCTTTTATTACCATATAGCTATTGGATTGGCCATCCGGT), in a 20µl total reaction. qPCR was conducted with an initial denaturation of 94**°**C for 10 min followed by 40 cycles of 94**°**C for 15 seconds and 60**°**C for 1 min. Inhibition was evaluated by comparing the Ct of eDNA spiked and unspiked reactions. An increase of greater than one Ct between spiked and unspiked reactions was assumed to indicate sample inhibition. As no samples showed evidence for inhibition, metabarcoding proceeded using the raw extract as follows.

Metabarcoding targeted a variable length (90-130 base-pairs without primer regions) section (V9) of the eukaryotic nuclear 18S ribosomal RNA that targets eukaryotes and variably amplifies some bacteria and archaea [45,46]. A qPCR with a melting curve was performed to determine the optimal number of cycles for the PCRs [47]. The qPCR was run in 20 μL reactions, consisting of 2 μL DNA template from a representative subset of samples, 1x AmpliTaq Gold 360 Master Mix, 0.8 μM forward and reverse primer, and 1 μl of SYBR Green/ROX solution (one part SYBR Green I nucleic acid gel stain (#S7563, Invitrogen, Waltham, MA, USA), four parts ROX Reference Dye (#12223-012, Invitrogen) and 2000 parts high-grade DMSO). The qPCR was conducted with an initial denaturation of 95**°**C for 10 min followed by 45 cycles of 95**°**C for 30 sec, 57**°**C for 30 sec and 72**°**C for 1 min followed by a terminal melting curve. To avoid overamplification, while still maintaining sufficient DNA for library construction, the number of cycles for the subsequent metabarcoding was selected by visual examination of when the majority of samples had reached the log-linear amplification phase. PCRs for eDNA metabarcoding were performed identically to the above qPCR with eight independently processed replicates with PCR 30 cycles, excluding the SYBR-ROX mix, and incorporating a final PCR extension of 72**°**C for 7 min in place of the melting-curve. The PCR forward and reverse primers incorporated unique dual nucleotide tags on the 5’-end such that each tag is only used in a single unique combined pair per sequencing library build (8 nucleotides in length, >3 differences between tags). Negative extraction controls and negative PCR controls (2-3 negative controls in every 40 PCRs) were amplified and sequenced alongside experimental samples. Following the PCR amplification, amplicons were visualised on a 2% agarose gel and equimolarly pooled according to the strength of the target DNA band (15 μL for very light/no band, 10 μL for medium strength band and 5 μL for strong band), 15ul of PCR product from extraction negative controls and PCR negative controls were added.

PCR replicates were distributed across pools using different tags for each PCR replicate and pooling generated a total of 24 amplicon pools, which were purified using MagBio HighPrep PCR clean-up system (MagBio Genomics, Inc, Gaithersburg, MD, USA) using the manufacturer-provided protocol with a 1:1 bead to amplicon pool ratio. To minimise tag-jumps, amplicon pools were built into libraries using the Tagsteady library protocol [48] with unique-dual indexes (different index for each forward and reverse adaptor, each index used only once) of 10 nucleotides in length. One library negative control was included in the library build and sequenced alongside experimental libraries. Libraries were cleaned up using MagBio HighPrep PCR clean-up system using the manufacturer-provided protocol with a 0.8:1 beads to library ratio. The amplicon libraries were sequenced using an Illumina NovaSeq 6000 instrument over over three SP flowcell (250 bp paired-end) lanes with a 10% PhiX library spike-in, for ca. 170,000 reads per individual PCR. Samples were sequenced alongside other libraries containing amplicons from different targeted regions.

### d) Metabarcoding bioinformatics

Illumina demultiplexed library build pools were further demultiplexed into PCR replicates using Cutadapt (v4.2) [49] by matching the primer and tag sequences for the forward and reverse primer in the read 1 and read 2 file, respectively, with only a single mismatch per primer allowed. This process was repeated on each pooled file, as the library build method used here creates libraries with the amplicon in both orientations, with the forward and reverse primer trimmed in the read 2 and read 1 files, respectively. Each orientation was then independently processed using DADA2 (v1.29.0) [50] in R (v.4.2.2) [51] under default settings unless detailed here. Demultiplexed reads were filtered using the *filterAndTrim* function with following parameters ‘maxN=0, maxEE=c(1,1), truncQ=2’. The NovaSeq instrument bins per base-pair sequence quality data, resulting in poor error model fitting using the *LearnErrors* function, therefore an error model enforcing monotonicity was used as outlined in the supplied bioinformatic script. Following ASV generation and chimera removal, the resultant data tables from each demultiplexing orientation were combined, summing ASVs with 100% identity (once oriented in the reverse complement). The following quality control filters were applied for each sequencing lane independently.

Singleton observations and ASVs observed only in a single PCR replicate per lane were discarded. Any observations from sediment samples with greater than the average number of reads per ASV across negative controls in the sequenced lane were set to zero. ASVs greater than 170 base-pairs or less than 70 base-pairs were removed. Datasets from each sequencing lane were then combined, combining observations for ASVs with 100% identity.

ASVs were taxonomically assigned using blastn (v.2.12.0+) [52] against the NCBI nt database (downloaded 27 Jan 2022) to return 200 hits (*-num_alignments 200*) per ASV; these were then parsed using a custom R script that uses a lowest common ancestor approach to assign a taxonomy to each ASV (ParseTaxonomy, doi:10.5281/zenodo.4671710). The *assignTaxonomy* function from DADA2 [53] with the protist-oriented PR2 database (v.5.0.0) [54] was used to generate a second set of assignments as this database uses a different taxonomic classification system and provides high-confidence higher taxonomic assignment (e.g. protist / metazoa). Finally, ASVs were assigned to functionally annotated sequences for the V9 region of 18S from PR2 database (DOI: 10.5281/zenodo.3768950) using vsearch (v2.28.1) [55] (sequence identity >85%, coverage >90%), with assignments only made in the case of consensus agreement among all reference sequence hits.

### e) Climatic and human context

Five previously published local climatic records were sourced to represent climatic changes concurrent with the newly collected eDNA data. Two annually resolved sclerochronological isotope records (δ^13^C & δ^18^O) which were derived from the shells of marine bivalve *Arctica islandica* sampled from the north Icelandic shelf [56,57]. The remaining three records were all derived from a sediment record (MD99-2275, 66.551667 -17.699833) collected in 1999 within 100m of the sampling site of PC19 presented here. These three records consisted of two sea surface temperature proxies, a diatom transfer function [58] and C_37_ alkenones [59], and one proxy for Arctic sea ice (IP25) [60]. 100-year splines were generated using the *detrend.series* function from the *dplR* package (v1.7.6.) [61] in R.

Icelandic human population estimates were sourced from reconstructions based on historical sources and modern government records [62–64].

To provide a broad, non-exhaustive (for example see [25]), centennial-scale proxy for human marine resource utilisation across Icelandic settlement history, 21 previously published archaeofaunal collections from household midden contexts across North and Western Iceland were sourced for marine fish and terrestrial mammal NISPs (Number of Identified Specimens). The midden remains are associated with NABO (North Atlantic Biocultural Organization) projects and comprise 34 period-specific animal bone collections in total [26,65–67]. See Supplementary File 1 for raw data and Supplementary Information 1 for a labelled map of sites. To ensure that poorly constrained assemblages did not have a disproportionate influence, the following measures were taken: for each century from the settlement of Iceland a weighted average proportion of marine and terrestrial bone specimens was calculated from all assemblages in the century, the weight for each assemblage was calculated by one divided by the number of centuries a given assemblage covers. For example, an assemblage constrained to two centuries would have a weight of 0.5 given in the weighted mean in each century (Supplementary File 1). We acknowledge that these temporal constraints remove any interpretative value based on archaeological context and that they therefore can merely provide a generalised overview across time.

### f) Statistical analysis

Taxonomic overview barplots were generated from datasets with the mean relative proportion per ASV across all eight replicates. Any ASV with less than 1% of reads was categorised as ‘other’ within the sample. Taxonomic groups from the functionally annotated PR2 assignments were used (10.5281/zenodo.3768950) for visualisation; assignments generated with *assignTaxonomy* in DADA2 were also used to generate visualisations at family level to confirm the annotated dataset.

ASV richness was estimated per sample by calculating the number of ASVs with non-zero values from mean relative proportion datasets calculated across all eight PCR replicates. Non-linear generalised additive models (GAMs) were generated using the *gam* function from the *mgcv* package (v1.9-0) [68] in R with k = 20 and using the restricted maximum likelihood method. Data subsets for metazoa, protists and bacteria were generated using domain level PR2 assignments at 80% bootstrap support from the *assignTaxonomy* function in DADA2. All ASVs assigned to Actinopterygii using the NCBI approach were checked manually against the online NCBI blast portal (last accessed Jan 2024). Following the assignment approach for cod and herring outlined in Holman et al. [14], ASVs were combined within individual PCR replicates if assigned identical genera (*Gadus* for cod, *Clupea* for herring). The number of positive PCR replicates was then calculated for each sample and the non-linear fit and 95% confidence interval of PCR replicates over time were produced using GAMs as above (k = 20, method = restricted maximum likelihood).

The effect of sea surface temperature (SST) on fish detection proportion was evaluated using a linear regression in R (function *lm*) for cod and herring separately. The detection proportion was regressed against the mean of the two SST proxies outlined above.

## Results

### a) Chronology

All radiocarbon ages and associated errors are presented in Supplementary Information 2 and electron microprobe data are presented in Supplementary File 2. The two tephra horizons sampled from the PC19 core both exhibit andesitic to dacitic geochemical composition characteristic of the Hekla volcanic system [40]. Based on the regional tephrochronological framework [41], similarity in probe-derived geochemical composition (Supplementary File 2) and the alignment with PC19 radiocarbon ages, we suggest the the upper tephra layer correlates to Hekla 1300 (H1300). While the stratigraphically lower tephra layer exhibits similar chemistry to the Hekla volcano, the tephra composition is not typical of Hekla 3 (a widespread c. 3 ka marker layer, described from the region). We therefore leave the ash un-attributed and merely present the chemical composition (Supplementary File 2 and Supplementary Information 3). The upper tephra layer correlates to the historic marker layer Hekla 1300 CE (H1300; ∼ 650 cal. BP). The H1300 tephra is derived from an historic eruption which deposited ash in a northward trajectory for roughly 12 months [69,70].

Age-depth models combining the above radiocarbon dates and the tephra layer showed continuous deposition and fairly constant sedimentation rates as shown in Supplementary Information 4.

### b) Environmental DNA

A total of 204 eDNA samples were amplified with eight PCR replicates per sample, resulting in 368 replicates for GC01 and 1,264 replicates for PC19. Sequencing produced a total of 456.4 million demultiplexed reads across the experimental samples, with an average of 150,083 ± 84,787 for each individual PCR replicate remaining after DADA2 processing and filtering. The proportion of sequences filtered out during primer trimming, DADA2 processing and subsequent quality control filtration was small (>25% per step - see Supplementary Information 5 for values). A total of 320 negative controls were sequenced producing a total of 48,237 reads.

Negative control samples had a mean of 151 ± 1512 (s.d.) reads per sample and a median of 4 ± 56 (IQR) reads per sample. A full overview of the negative controls and their taxonomic assignment is provided in Supplementary Information 6. ASV generation and filtering produced a final dataset for analysis with 20,453 ASVs for GC01 and 24,160 ASVs for PC19 with a combined total of 31,831 ASVs of which 12,782 ASVs were detected in both cores.

Using the PR2 database, taxonomic domain could be assigned to 24,381 of the 31,831 ASVs with 80% bootstrap support across both cores. Of these, 22,659 of the ASVs were assigned to Eukaryota, 1701 to Bacteria and 21 to Archaea. From the Eukaryotic ASVs, 137 classes, 269 orders, 512 families and 901 genera could be assigned with 80% bootstrap support. High quality taxonomic assignments (>99% identity, >90% coverage) with the larger, but relatively less curated, NCBI database could be made for 2,606 ASVs; from these ASVs 731 genera could be annotated.

Overall, when considering the assemblage composition at a high taxonomic resolution, (Fig. 1) the ecosystem displayed relative stability across the time series. Both cores showed similar patterns in the taxonomically assigned fraction, with a greater proportion of assigned reads in the PC19 compared to GC01.

**Figure 1.**
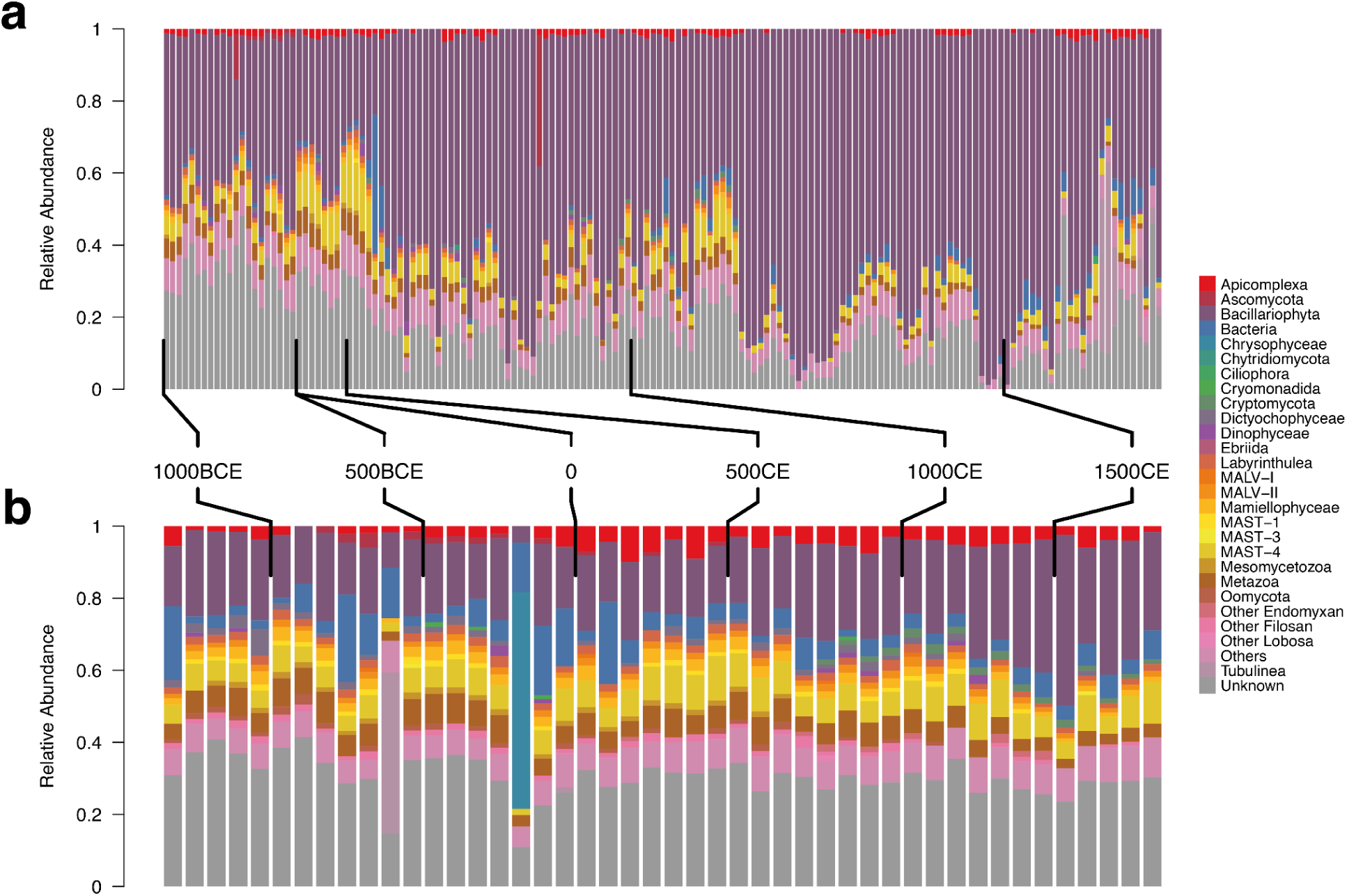
Barplots showing proportion of sequences assigned to taxonomic groupings from metabarcoding of marine sediments collected from the north Icelandic shelf. Calendar age brackets for samples are indicated by ticks for **a)** PC19 and **b)** GC01 core. Colours indicate broad taxonomic grouping with the group per colour provided in the right hand legend.

Across almost all samples the largest proportion of assignable reads corresponded with *Bacillariophyta* (Diatoms), as shown in purple in Figure 1. A single ASV contributed a substantial proportion of diatom sequences, with over 132 million sequences across all samples. This ASV was assigned to the genus *Chaetoceros*, a common diatom genus that frequently dominates Icelandic spring phytoplankton assemblages [30]. Across both cores the proportion of sequences assigned to this ASV increased towards the present day as shown in Supplementary Information 7.

ASV richness patterns differed by taxonomic group with bacteria showing a complex topology with little clear overall trend across the study period apart from an increase in richness in both cores across the final 200 years of the series (see Supplementary Information 8). Protist and animal richness was more stable in comparison, with greater variability in the series in the high-resolution PC19 core (see Supplementary Information 8). For both protist and animal subsets, GC01 showed approximately 30% lower richness in the oldest samples. ASV richness converged in the most recent 300 years of the time series for both taxonomic subsets (see Supplementary Information 8).

Two ASVs were taxonomically assigned to the Genus *Gadus* (cod) and six to *Clupea* (herring). The resultant proportion of positive detections for cod and herring are shown in Figure 2a & b below. Overall, both cores showed no clear change in the detection proportion of cod. In contrast, herring detections showed a clear decrease across the entire time series across both cores with samples showing no detection of herring only in the most recent millenia.

**Figure 2.**
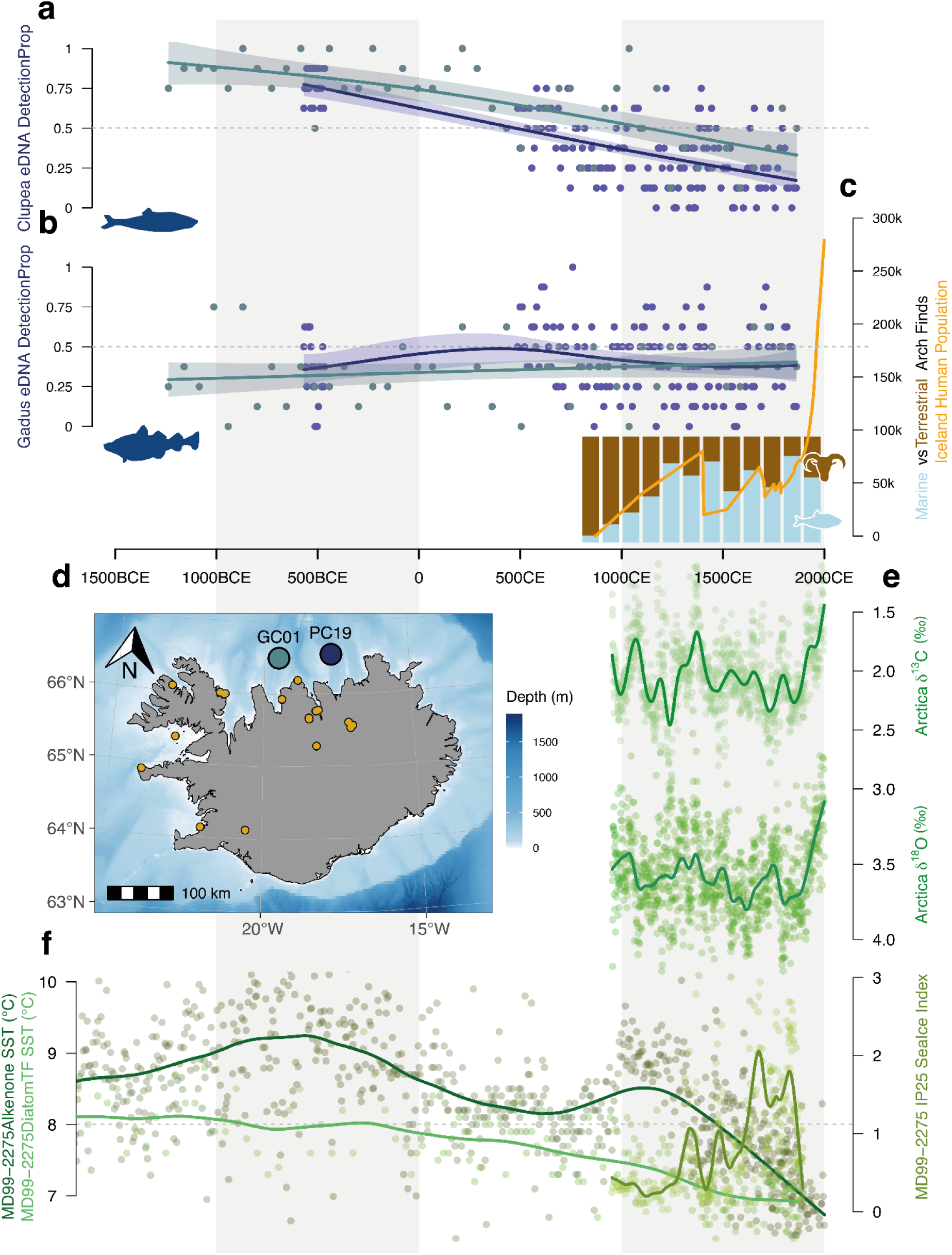
Proportion of positive PCR replicates for ASVs assigned to **a** Herring (Genus Clupea) and **b** Cod (Genus Gadus) over time for PC19 (navy points) and GC01 (teal points). Generalised additive model fits are shown with 95% confidence intervals for PC19 (teal line) and GC01 (navy line) and dashed grey lines indicate 0.5 detection proportion. **c** Icelandic human population(orange line) and average proportion of marine vs terrestrial zooarchaeological finds across Iceland. **d** Map of Iceland showing core sampling locations for GC01 (teal) and PC19 (navy) and archaeological sites used to calculate proportion of marine vs terrestrial finds. **e** Annually resolved Arctica islandica δ^13^C and δ^18^O shell series plotted with a 100-year loess first-order low-pass filter (green lines). **f** Sediment core climate proxies with 100-year loess first-order low-pass filter for SST derived from alkenone (dark green) and diatom transfer function (light green), IP25 sea ice proxy shown in olive green (right hand axis).

### c) Environmental and human context

Figure 2c shows the Icelandic human population increasing rapidly over several hundred years after settlement in the late 9th century CE, reaching a documented peak in 1400 CE and subsequent decline over only five years as a result of the Black Death [62]. The population remained below the pre-plague peak, experiencing slow recovery and a second fast decline because of smallpox in the 1700s, and only reaching the size before the Black Death in the 1900s.

Alongside these population changes, Icelandic zooarchaeological remains revealed two broad marine versus terrestrial resource use patterns (Figure 2c). Firstly, site profiles showed an increase from a predominantly terrestrial mammal focus to a marine fish dominated focus across the 9th-13th century. Secondly, the 14th-20th century site proportions suggested a relatively stable focus on marine fish as a major resource (except the 16th century with 48.3% marine proportion of finds). These century-scale estimates show only the broadest patterns as context, and site specific, socio-economic nuances and taphonomic processes are not shown by these estimates.

The δ^13^C annually resolved record (Figure 2e) showed considerable multidecadal-scale variability, with no overall long-term trend, and a rapid decrease in the last century as a result of the marine ^13^C Suess effect. The δ^18^O record (Figure 2e) also showed variability, recovering previously described climatic periods such as the Medieval Climate Anomaly and the Little Ice Age [57], in addition to a rapid decrease in the last century driven by anthropogenic climate change. This terminal decrease aside, the overall trend in the δ^18^O record is a cooling climate punctuated by variability. This agrees with the sea surface temperature records (Figure 2f, left side axes) which both show a cooling trend across the entire period, notwithstanding two increases in the diatom transfer function record centred around 500 BCE and 1200 CE. Finally, the IP25 record (Figure 2f, right side axes) shows a punctuated increase across the last millenia, adding support for an overall cooling climate with increased sea ice.

### d) Cod and Herring SST Correlations

There was no significant linear relationship between cod detection proportion and sea surface temperature (p > 0.05, Figure 3a). However, there was a strong positive linear relationship between herring detection proportion and sea surface temperature (R^2^ = 0.49, F _3,200_ = 66.95, p < 0.001, Figure 3b) across both cores, with no significant difference between the slope of the cores (p = 0.957).

**Figure 3.**
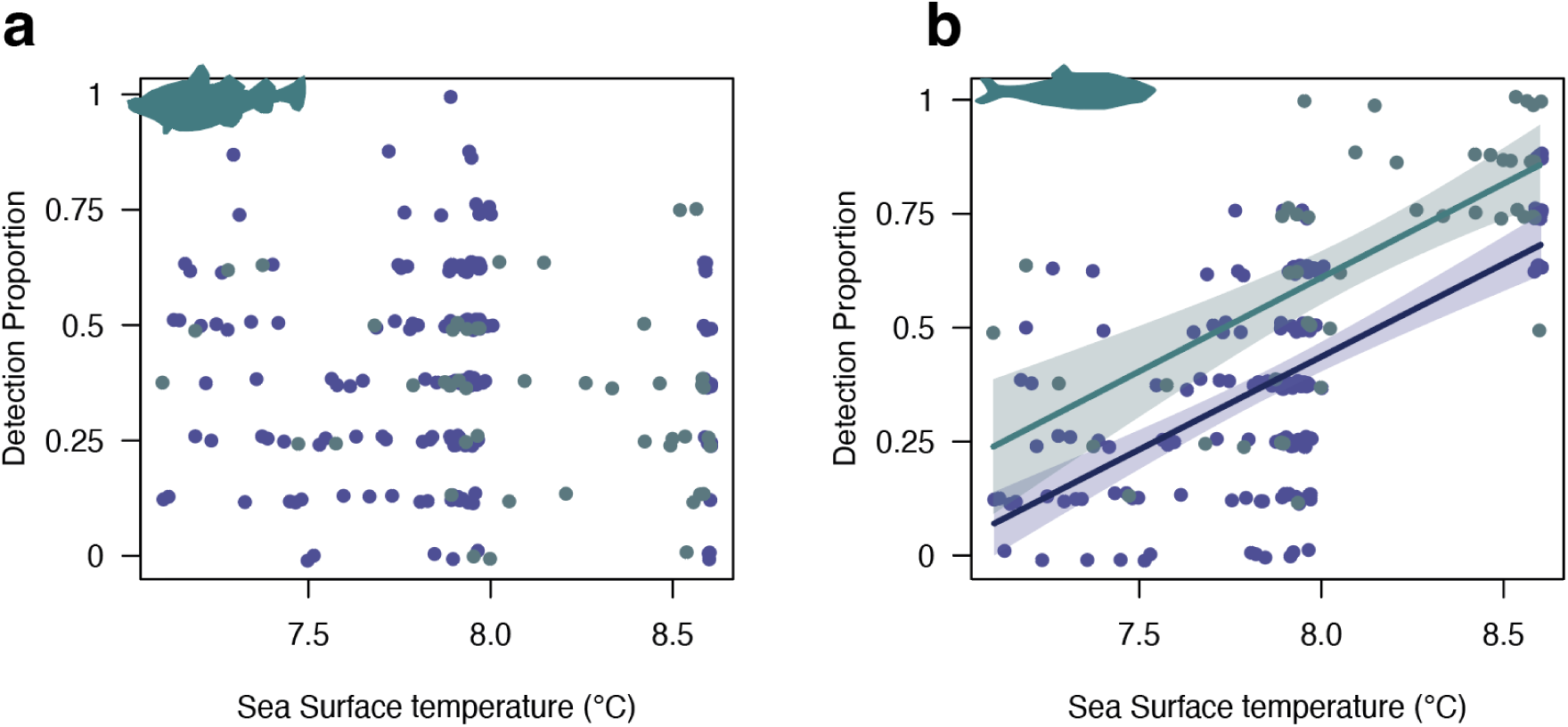
Proportion of positive PCR replicates for ASVs assigned to **a)** Cod (Genus Gadus) and **b)** Herring (Genus Clupea) against reconstructed sea surface temperature proxy estimates for two sediment cores; PC19 (navy points) and GC01 (teal points). The fit for a statistically significant linear regression model predicting herring detection from sea surface temperature is shown for PC19 (navy points) and GC01 (teal points) with the 95% confidence interval shown for each fit in shaded colours.

## Discussion

Natural climate change, human activities and biodiversity affect, and are affected by, one another in a complex nexus that can make it difficult to disentangle the relative contributions of each driver. For example, Rivera Rosas et al.[71] show a shift in biodiversity over a 15 year period from the Atacama deep-sea trench that correlated with both a strong El Niño climate oscillation event and extensive fishing in the area, leaving it unclear which of these drivers has the largest effect. In contrast, Ficetola et al.[72] revealed a limited role for natural climate when compared with the human-introduction of invasive rabbits on the sub-Antarctic Kerguelen Islands; the rabbits altered plant community structure and soil erosion rate, dramatically altering the terrestrial ecosystem. Here, we leverage the late settlement of humans on Iceland to determine whether climate or human activities have a greater effect on marine biodiversity.

Overall, we found limited anthropogenic impacts on marine biodiversity across the study period, with little turnover of broad taxonomic groups, or sharp changes in richness, related to human settlement. Furthermore, we find no evidence for changing detection frequencies of cod, which we infer as incidence through time, in response to fishing pressure. In contrast, we observe decreasing herring detection frequencies from 1500 BCE - 1700 CE, showing a strong positive correlation with sea surface temperature proxies. Overall, these results suggest that climate, rather than human activity, has been an important ecological force across the study period in northern Iceland.

Biodiversity change is multifaceted. Species richness, turnover and abundance all describe different features of biodiversity, which may in turn change depending on the scale at which they are assessed [73]. Considering the post-settlement population growth in Iceland (Figure 2c), widespread reliance on marine resources and exports (Figure 2c & Hambrecht et al.[26]), and broad human impact on Iceland’s terrestrial ecosystems [19], we might expect to find evidence of human-driven biodiversity changes in Icelandic seas. However, our data show limited changes in the relative abundance of broad taxonomic groups (Figure 1) related to human settlement. Previous work has shown that anthropogenic activities can leave substantial shifts in broad taxonomic groups in marine sediment records [12,74], but also that relative read abundance does not consistently reveal differences in different ecosystems [75].

Clearly preindustrial humans had some impact on Icelandic marine ecosystems, notably the local extirpation of walrus [32], but relative to documented examples of anthropogenic large-scale taxonomic turnover [12] or substantial shifts in richness [71] it appears human influences were relatively minor. Preindustrial Icelandic settlers are most likely to have affected marine biodiversity through fishing [3], as other principal drivers of biodiversity change (climate change, pollution, seascape use change) seen in modern ecosystems are typical of industrialisation [3]. We show no changes in the detection rate of cod, the most commonly exploited species [26], throughout the time-series (Figure 2b), in agreement with studies that show no overall human impact on cod trophic position, diet or population size [33–36]. Anthropogenic marine impacts in Europe on fish and fisheries can be seen as early as the Middle Ages, with sedimentation and land use change of anadromous fish habitats impacting on salmon and sturgeon populations [76]. The substantially lower population size of Iceland compared to Europe across the study period (Figure 2c), against the backdrop of highly productive oceans with substantial fish stocks [30,77], make it likely that preindustrial Icelandic fish populations were predominantly unaffected by human impact.

In contrast, natural climatic variation has played a key role in changing Icelandic ecosystems [26,30,78]. The north Icelandic shelf marine ecosystem is located at the confluence of the relatively cold, fresh polar water of the East Icelandic current and the more saline, warm Atlantic water of the Irminger current. These water bodies contend and mix at the Polar Front such that one current system typically dominates, resulting in fresher cooler periods and more saline warmer periods [30]. These periods may persist for several years, resulting in temperature changes in the North Icelandic shelf that can differ by as much as 4°C [30]. The decreasing SST proxies (Figure 2f) suggest increasing polar water influx across the study period, with correlating increases of sea ice (Figure 2f) characteristic of increased polar influx. The cooler polar water generally results in lower primary productivity as a result of early nutrient exhaustion during spring in polar water conditions [30], lower zooplankton growth rates at cooler temperatures, and lower zooplankton densities in response to decreased phytoplankton. Cumulatively this means years dominated by polar waters have lower zooplankton densities to support secondary consumers, such as herring and capelin (*Mallotus villosus*). Our data show decreasing detections of herring over time and a positive correlation between SST and herring detection rate, indicative of a larger proportion of years with high polar water input resulting in lower herring densities. We do not, however, see the same pattern for cod, despite capelin being a critically important prey item [30,79].

One important caveat in understanding past ecosystem dynamics is that our knowledge of a contemporary ecosystem may not reflect how the ecosystem worked in the past. Research has documented whole ecosystem shifts in response to anthropogenic [12] and natural stressors [11], and fish range shifts have been documented both in past ecosystems [80], and more recently in species over the last 100 years [77].

Assuming the north Icelandic shelf ecosystem has broadly similar structure in the past compared to today, we can explore potential explanations for why herring exhibit stronger climate responses than cod. There are several overlapping herring stocks in Iceland [81], two have feeding grounds on the north Icelandic shelf (Icelandic spring-spawning herring - ISPH Norwegian spring-spawning herring - NSPH) but vary in their spawning location and size, with the NSPH being much larger [31]. During colder conditions with high polar water input the NSPH stock can shift feeding grounds east, outside of Icelandic waters [31]. The increase in cooler years seen in our data may result in the NSPH feeding more frequently east of Iceland, which would produce the correlation between SST and herring detection rate observed here. Like the ISPH, Icelandic capelin breed in the southern coasts of Iceland and have similar environmental spawning requirements [77]. However, their feeding grounds are much more northerly (68° - 72°) [79], and thus may not respond to climatic variation on the north Icelandic shelf in the same way as herring. Nonetheless, evidence indicates that there is a relationship between mean Icelandic capelin weight and northern Icelandic environmental conditions [79], but the same analysis showed no relationship between zooplankton biomass on the north Icelandic shelf and capelin weight.

While capelin are an important food source for cod [30,79], the stability in cod detection rates may be explained by cod shifting their feeding to other taxa that are less affected by natural climatic variation. Evidence from gut-content-analyses of modern Icelandic cod indicate that, particularly for smaller cod, there are many different prey taxa on which cod populations can feed [82]. Environmental DNA quantity is more representative of organismal biomass than individual counts [10,83], therefore the observed stability in cod detection rates could encompass a range of populations with cod of different size possibly driven by changing availability of prey. Another explanation, and a common limitation of most sedaDNA studies, is the representativeness of a small number of sediment records of wider ecological patterns. It is possible that the observed cod detection rate patterns, among the other detected biodiversity patterns, reflect only a limited picture of the entire changing ecosystem. We currently lack an understanding of the ecosystem area or “catchment” represented by a sediment record. Studies using modern eDNA have shown that taxa can be detected over 1000m from their source [84], but also that distinct marine communities can be delimited over less than 100m [85]. Given the consistent patterns between cores (Figure 1 & 2, Supplementary Figure 9), we can be confident that the sediment records presented here are broadly representative of the dominant ecological changes in the study system.

The consistency of biodiversity patterns observed between the cores is also encouraging considering they were collected over two decades apart. Previous work in lake sediments indicates that there may be changes in biodiversity, particularly in prokaryotic communities, during storage at 4°C that can add bias to the relative abundance of reads from sediment samples [86]. However, work in marine sedimentary DNA suggests that these changes may be minor and, provided the sediment has been stored appropriately, sedimentary DNA can reconstruct broad community shifts and species incidence [14,87]. Here we observe similar overall taxonomic patterns between the cores (Figure 1), albeit with a greater proportion of unassigned reads in the older core, richness and detection patterns between the two cores. The cores even show very similar detection rates for taxa with complex incidence patterns over time (Supplementary Information 9), providing confidence in the methodology across the workflow.

While the records presented in this work end in the pre-industrial period, the narrative of Icelandic marine exploitation has continued. The industrialisation of fisheries and human-driven climate change has had a dramatic effect on marine ecosystems [3], resulting in large changes in fish populations, with shifts in trophic patterns [82] and even complete collapse of some Icelandic stocks [31,81]. Future work on eDNA data from sediment records extending into the industrial period would help link the observed patterns here to modern datasets, allowing a complete record from the present day, documenting marine biodiversity change across all of Icelandic human history.

## Acknowledgments

We thank Árni Daníel Júlíusson for valuable discussions and assistance with Icelandic human population data. Special thanks to the GeoGenetics Sequencing Core at the University of Copenhagen for their essential work and support in DNA sequencing. We also acknowledge and thank the National Marine Facilities staff and the RRS Discovery crew, for their involvement in the 2022 DY150 cruise. We thank Bent Petersen for his continued support with the Mjölnir high-performance computing cluster. Additionally, we acknowledge the staff at the SUERC AMS laboratory in East Kilbride, Scotland, and Dr. Xiaomei Xu at the Keck Carbon Cycle AMS Facility, University of California, Irvine, for their expertise in AMS 14C dating. Finally, LH extends gratitude to the Arctic Circle for the support provided through the Grímsson Fellowship, which allowed this work to be completed during a stay in Iceland.

## Data, code and materials

All raw sequencing data has been uploaded under European Nucleotide Archive accession PRJEB78865. All code, analysis outputs and intermediate datasets used in the generation of results have been permanently archived at Zenodo under DOI: 10.5281/zenodo.13758187.

