## Supplementary Information for "Ancient environmental DNA indicates limited human impact on marine biodiversity in pre-industrial Iceland"

**Supplementary Information 1:** Map of Iceland showing named archaeological sites (yellow points) used for reconstruction of marine versus terrestrial exploitation.

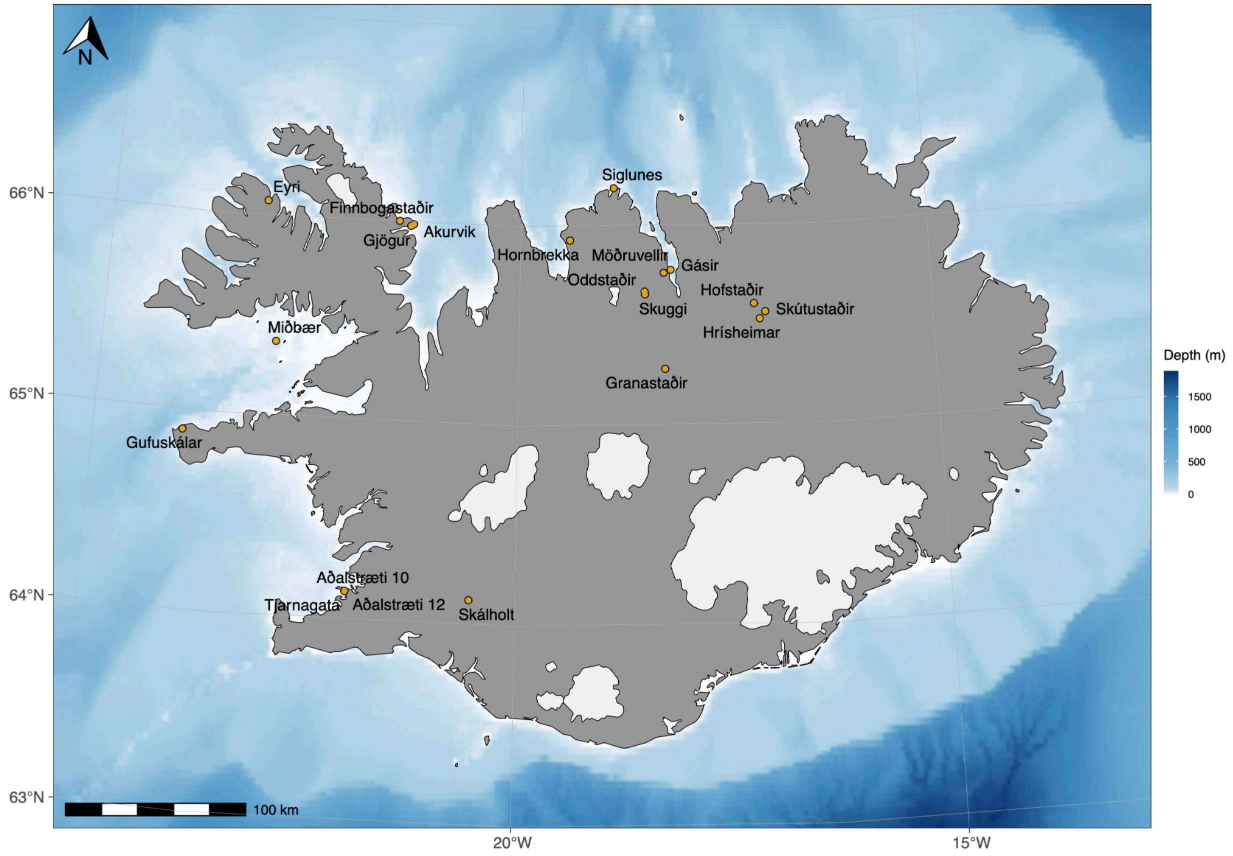

**Supplementary Information 2 :** Radiocarbon ages generated for sediment cores GC01 and PC19.

| Identifier | C14 age | C14 age±1sd | depth_mean | depth_range | core | Percent modern carbon (%) | Percent modern carbon (%)±1sd | carbon_% |
|---|---|---|---|---|---|---|---|---|
| UCIAMS-284082 | 1847 | 35 | 48.5 | 48-49 | GC01 | 79.46 | 0.35 | 7.38 |
| UCIAMS-284081 | 2210 | 35 | 80.5 | 80-81 | GC01 | 75.94 | 0.33 | 8.93 |
| UCIAMS-275845 | 3668 | 35 | 176.5 | 176-177 | GC01 | 63.34 | 0.28 | 11.44 |
| UCIAMS-275837 | 705 | 35 | 15 | 14-16 | PC19 | 91.6 | 0.4 | 11.44 |
| UCIAMS-284084 | 1277 | 35 | 109 | 108-110 | PC19 | 85.31 | 0.37 | 10.67 |
| UCIAMS-284085 | 1756 | 35 | 199.5 | 199-200 | PC19 | 80.37 | 0.35 | 11.37 |
| UCIAMS-275838 | 2327 | 35 | 275.5 | 275-276 | PC19 | 74.85 | 0.33 | 11.63 |
| UCIAMS-284086 | 2693 | 35 | 339 | 338-340 | PC19 | 71.52 | 0.31 | 11.35 |
| UCIAMS-275839 | 2994 | 35 | 400.5 | 400-401 | PC19 | 68.89 | 0.3 | 10.81 |
| UCIAMS-284087 | 3192 | 35 | 449 | 448-450 | PC19 | 67.21 | 0.29 | 10.4 |

**Supplementary Information 3 :** Comparison of CaO/K20 and FeO/TiO2 ratios of PC019 tephra (orange) to tephra marker layers from the Hekla volcano, previously described from the Iceland shelf (MD99; Larsen et al. 2002; Eiriksson et al. 2004). Plots A) and B) exhibit the similarity between PC019:109.5 and the Hekla 1300 marker layer (H1300). Plots C) and D) exhibit contrast between PC019:450 tephra and Hekla 3.

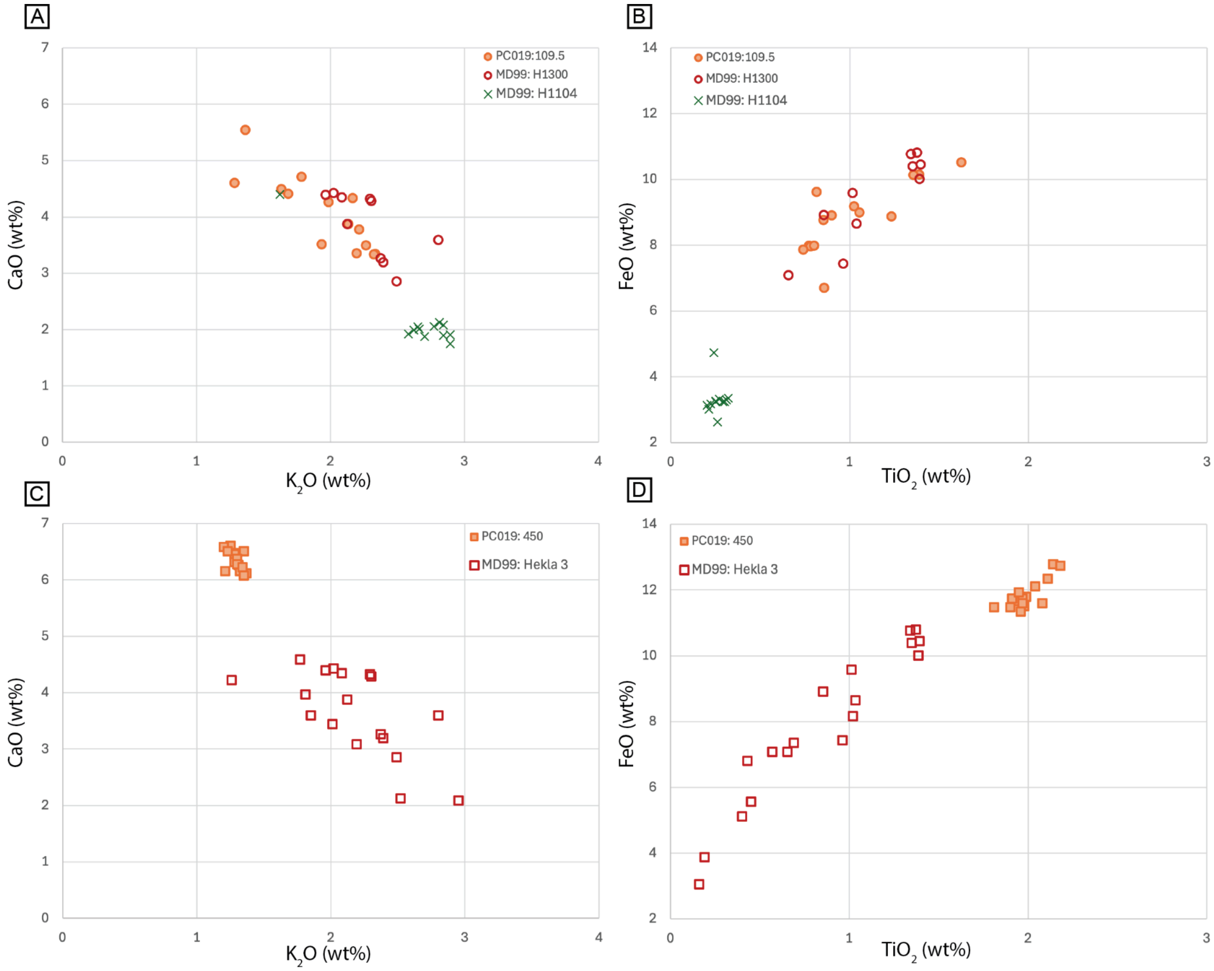

**Supplementary Information 4 :** Age-depth model outputs from rBacon for GC01 (left) and PC19 (right). For each age-depth model the diagnostic plots are shown in the top row and the age-depth model on the bottom. C14 ages are shown with probability distributions in blue, tephra dates are shown with turquoise vertical lines. Samples used for sedaDNA analysis are shown with yellow points. The dashed grey lines indicate the 95% model confidence intervals and the red dashed line the mean model.

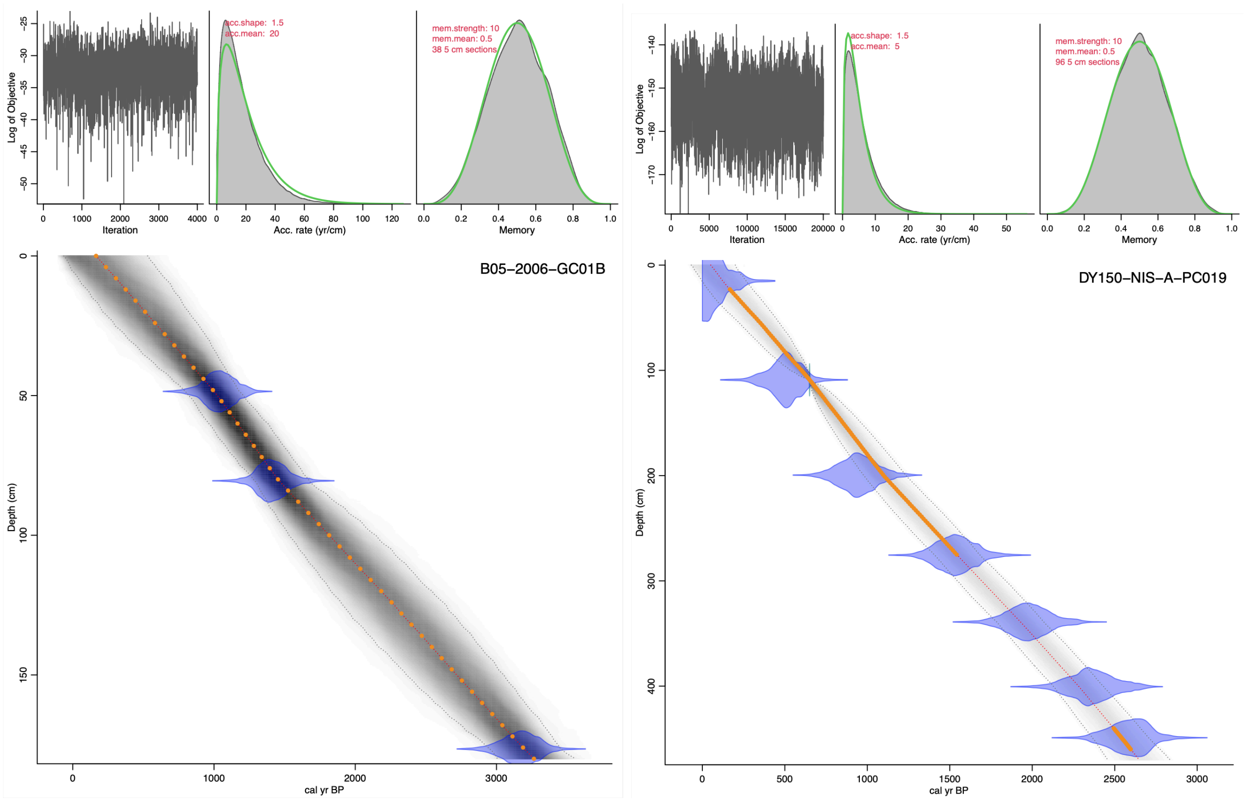

**Supplementary Information 5 :** Overview of raw and processed sequencing counts

| | ExperimentalMean | ExperimentalSD | NegativeControlMean | NegativeControlSD | SummedTotalExperimental | SummedTotalNegative |
|---|---|---|---|---|---|---|
| Initial Demultiplex | 197445.389 | 132201.792 | 181.596875 | 1786.69951 | 456493739 | 58111 |
| RevPrimerStripped | 194218.218 | 130505.92 | 177.803125 | 1756.37884 | 449032520 | 56897 |
| PostDADA2 | 150082.626 | 84787.4581 | 150.740625 | 1508.89476 | 244934846 | 48237 |

**Supplementary Information 6 :** Negative control overview

Negative controls produced a total of 48,237 reads across 320 samples. From these samples, extraction negative controls produced 19,780 reads and PCR controls produced 28,457 reads. Across all negative controls these reads generated 455 ASVs, taxonomic assignment (80% bootstrap agreement to the PR2 database as main manuscript) of these ASVs gave 321 eukaryotic 73 prokaryotic and 6 archaeal assignments. Negative control ASVs could be assigned to a total of 51 classes, 74 orders, 86 families and 125 genera.

Controls had a mean of 151 ± 1512 (s.d.) reads and a median of 4 ± 56 (IQR) reads. Controls had a mean of 2.5 ± 16.0 (s.d.) ASVs and a median of 1±2 (IQR) ASVs.

The below figure shows the taxonomic assignment at **a** subdivision level **b** family level for reads found in negative controls from DNA extraction and PCR.

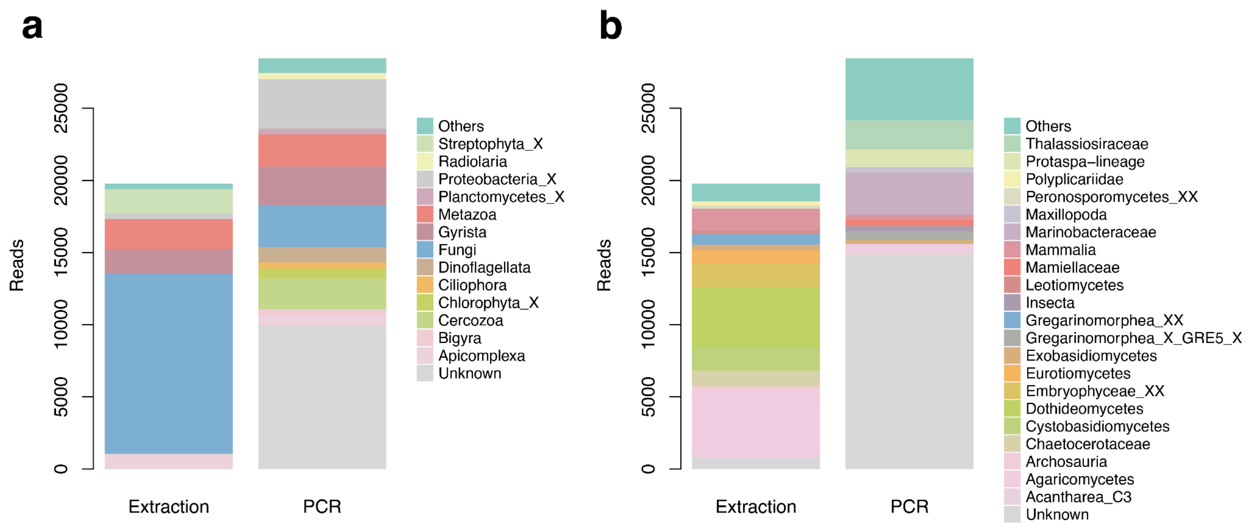


A large proportion of the reads in the negative samples came from fungi and unassigned ASVs. This is broadly the expected taxonomic diversity of control samples as extraction kits can contain fungal contaminants and PCR amplification of negative samples can result in amplification of non-informative PCR artefacts [1–4]. However, we also observe ASVs assigned to taxa found in the experimental samples. This too is expected, the DNA library preparation method used here minimises, but does not entirely remove tag jumps [5]. We can therefore expect a small amount of bleed over between samples, which is then removed bioinformatically as detailed in the main manuscript. Across all negatives 1,218 reads could be assigned to genus *Chaetoceros,* suggesting a bleed across between samples of <0.001%.

**Supplementary Information 7:** Proportion of total filtered reads per sample from the most frequently observed ASV (assigned genus Chaetoceros) across PC19 (navy) and GC01 (teal) sediment cores.

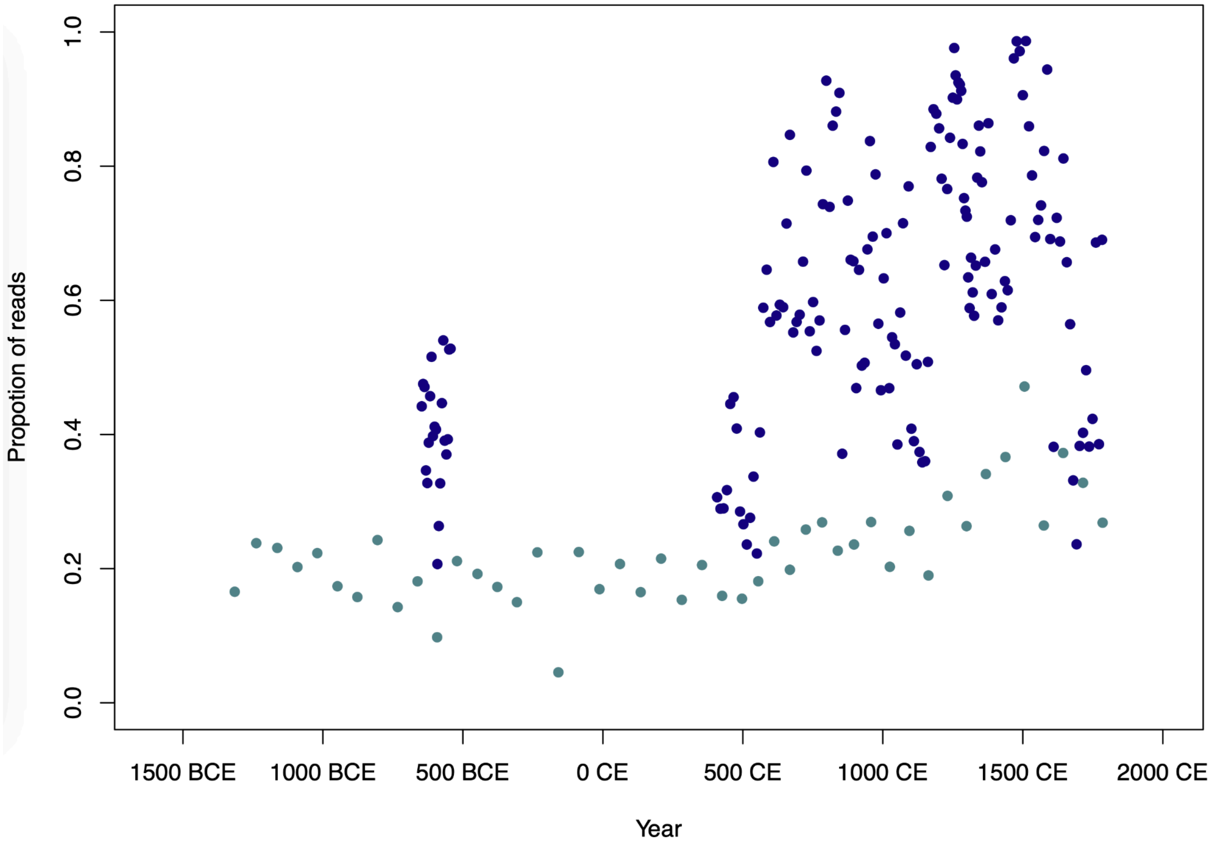

**Supplementary Information 8:** ASV Richness patterns over time for PC19 (navy points) and GC01 (teal points). Generalised additive model fits are shown with 95% confidence intervals for PC19 (teal line) and GC01 (navy line).

*Bacteria ASV richness*

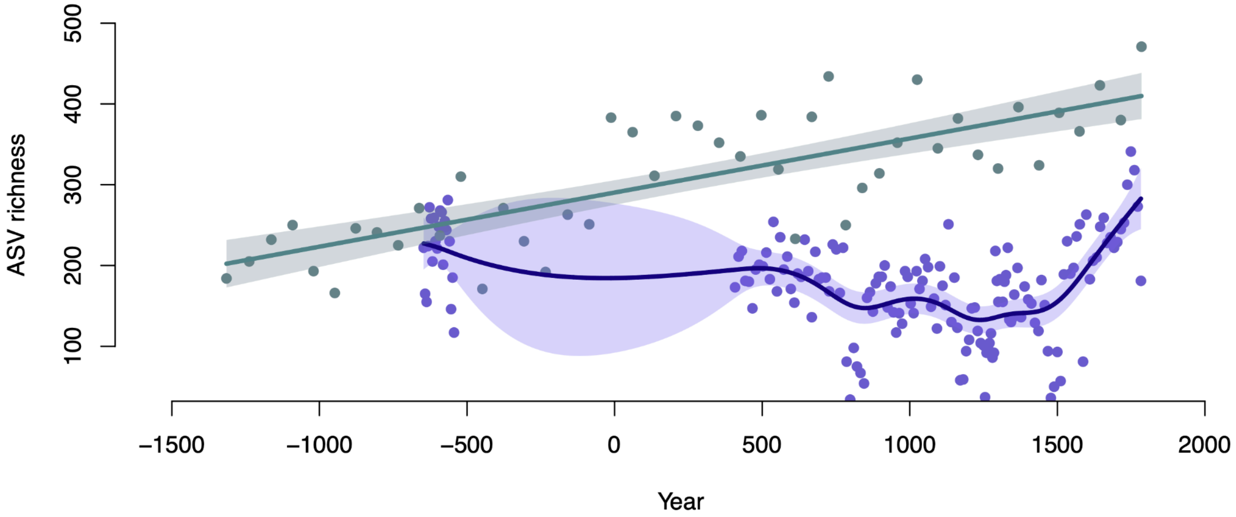


*Protist ASV richness*

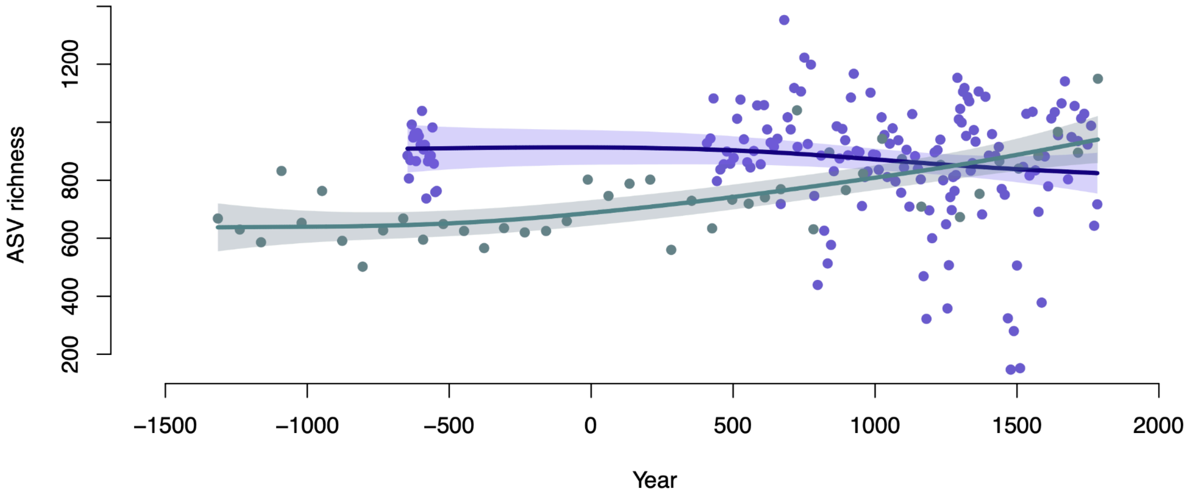


*Metazoan ASV richness*

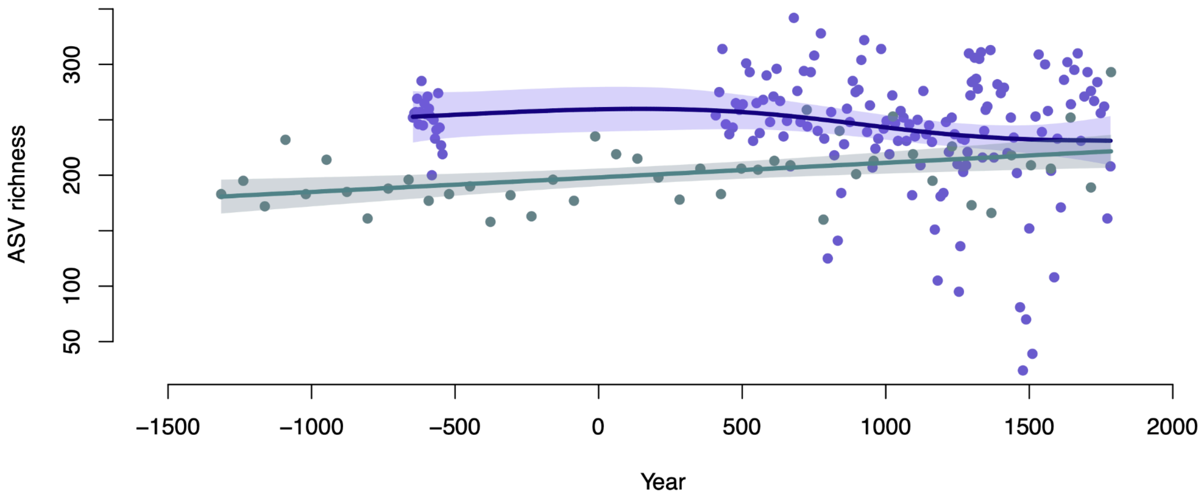


*All ASV richness*

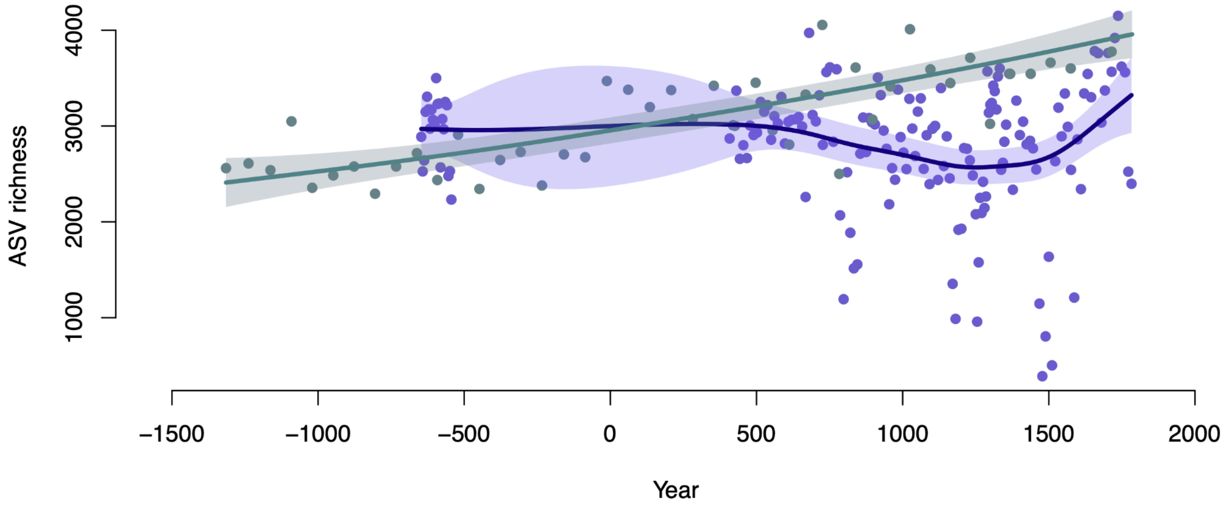

**Supplementary Information 9 :** Proportion of positive PCR replicates for ASVs assigned to the Helmet Jellyfish (Periphylla periphylla) over time for PC19 (navy points) and GC01 (teal points). Generalised additive model fits are shown with 95% confidence intervals for PC19 (teal line) and GC01 (navy line) and dashed grey lines indicate 0.5 detection proportion.

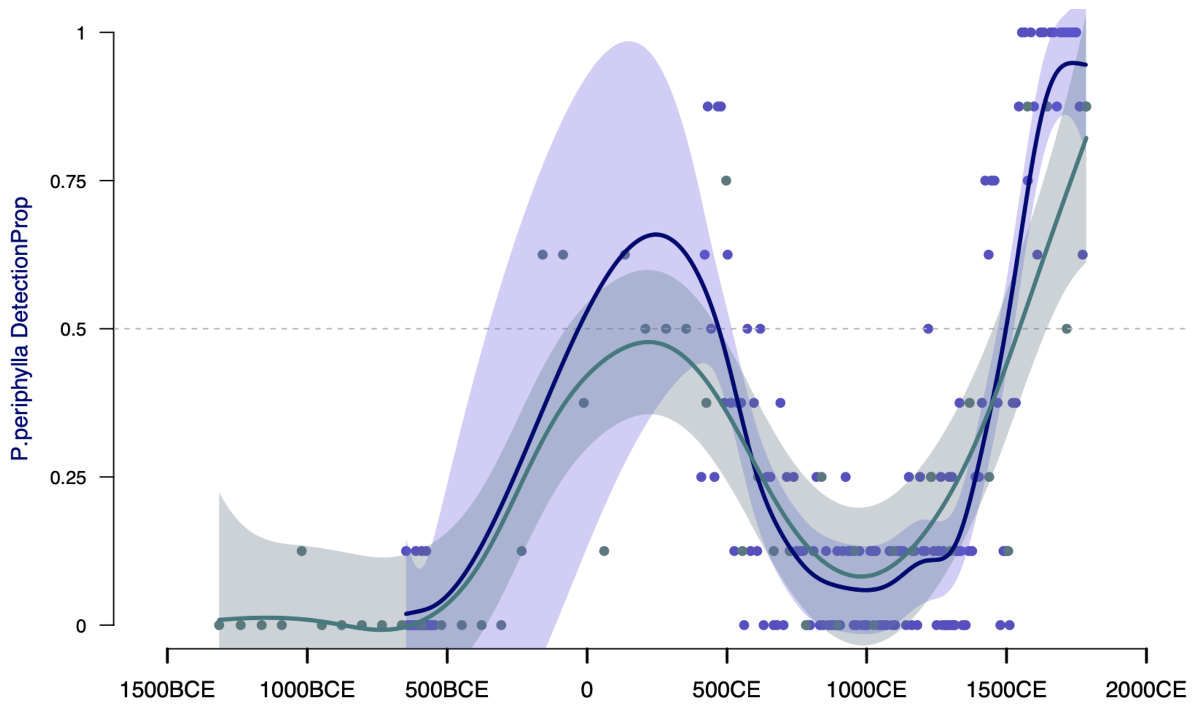
